# The molecular landscape and microenvironment of salivary duct carcinoma reveal new therapeutic opportunities

**DOI:** 10.1101/810028

**Authors:** Melissa Alame, Emmanuel Cornillot, Valère Cacheux, Guillaume Tosato, Marion Four, Laura De Oliveira, Stéphanie Gofflot, Philippe Delvenne, Evgenia Turtoi, Simon Cabello-Aguilar, Masahiko Nishiyama, Andrei Turtoi, Valérie Costes-Martineau, Jacques Colinge

## Abstract

Salivary duct carcinoma (SDC) is a rare and aggressive salivary gland cancer subtype with poor prognosis. The mutational landscape of SDC has been described rather exhaustively; yet, with respect to functional genomics and tumor microenvironment, little is known. In this study, transcriptomics and proteomics were combined to obtain the first characterization of the pathways deregulated in SDC. The data revealed the importance of Notch, TGB-β, and interferon-γ signaling. After associating computational biology, immunohistochemistry, multiplexed immunofluorescence, and digital imaging the first steps towards charting the cellular network within the microenvironment was initiated. According to immune infiltrate, two well-defined groups of tumors were observed, novel SDC immune checkpoints were discovered, and the key role played by macrophages and potentially NK cells in immunosuppression was shown. Furthermore, a clear trend between recurrence-free survival and M2 macrophage abundance was apparent. Independently, a measure of desmoplastic stromal reaction as determined by α-SMA abundance, was also shown. Altogether, these many findings open new perspectives for understanding and treating SDC. Before applying an immunotherapy, classical patient stratification according to immune infiltrate should be taken into account. Moreover, the microenvironment offers new potential targets including macrophages or NK cells, or even fibroblasts or hyaluronic acid. Related therapies that have been developed against, *e.g.*, pancreatic tumors could inspire equivalent therapy for SDC.

**Additional information:**

- Financial support: MA (1 grant, GIRCI SOOM API-K 2016-811-DRC-AC), JC (2 grants, Fondation ARC PJA 20141201975, Labex EpiGenMed ANR 10-LABX-0012), AT (2 grants, Gunma University GIAR Research Program for Omics-Based Medical Science, Labex MabImprove ANR 10-LABX-0053 starting grant), ET (1 grant, SIRIC Montpellier Cancer Grant INCa_Inserm_DGOS_12553).
- No conflict of interest
- 5408 words, 1 table, and 4 figures

**Statement of translational relevance:** Based on the presence or absence of an immune infiltrate, two groups of SDC were identified. These have the potential to provide a rationale for therapy and clinical trial enrolment. Two novel immune checkpoints that could be targeted were also identified; namely, CTLA-4/CD86 and TIM-3/galectin-9. Both showed the important contribution that macrophages and NK cells have in immunosuppression. Treatments that induce reprogramming or elimination of these cells could be considered. Moreover, the importance of the desmoplastic stroma was stressed. The stroma acts as a physical barrier against therapy suggesting that strategies developed against pancreatic tumors could inspire SDC treatments. For SDC devoid of immune infiltrate, components of the stroma including fibroblasts or hyaluronic acid could be targeted, *e.g.*, in combination with drugs against immune checkpoints or mutated genes. Finally, evidence that Notch and TGF-β signaling are prevalent in SDC was obtained. This translates into additional therapeutic options.

## Introduction

According to the World Health Organization (WHO) classification (1), salivary gland tumors constitute a heterogeneous group of tumors comprised of 24 histotypes. Among these, salivary duct carcinoma (SDC) represents 2% of the epithelial primary salivary gland tumors. It is an uncommon entity that predominantly arises at the parotid or submandibular region and is characterized by the degree of aggressiveness and the high mortality rate. At diagnosis, patients often present with lymph node involvement. Standard therapy primarily relies on local resection with adjuvant radiotherapy. Nevertheless, recurrences and distant metastases frequently occur. These respond poorly to chemotherapy; the usual second line treatment (2). Due to the associated histology, androgen receptor (AR) and human epidermal growth factor receptor 2 (HER2) positive-staining; WHO classify the neoplastic tissue as resembling invasive ductal mammary carcinoma (IDC) (1). Initial studies have reported potential efficacy of androgen deprivation therapy (ADT) (3) or HER2 inhibition, *e.g.*, with trastuzumab (4). Unfortunately, no pre-clinical models of SDC are available.

Significant efforts have been devoted to survey SDC somatic mutations and identify genetic insults that can be targeted (5–11). These studies revealed mutations in cancer genes involved in DNA damage, mitogen-activated protein kinase signaling, receptor tyrosine kinase, PI3K-AKT signaling, androgen signaling, histone modifiers, and other categories. Recurrent-mutated genes are *TP53* (depending on the report, 31%-68% of patients), *PIK3CA* (9%-37%), *HRAS* (11%-27%)*, FOXA1* (0%-25%), and *NF1* (0%-18%). Some mutations may limit the benefit of ADT, *e.g.*, *FOXA1* (5). Gene copy number alteration (CNA) analysis has indicated a modest rate of chromosome arm amplifications or deletions (5,8) and a limited number of gene fusion events have been identified (5). The transcriptomes of 16 SDCs were obtained and exploited to show transcriptional resemblance with breast tumors. To date, however, no comparison of SDC *versus* normal adjacent salivary duct tissue has been conducted to discover SDC-deregulated pathways.

The contribution of the tumor microenvironment (TME) to tumor progression and therapy resistance is major with most solid tumors (12). Immunotherapies have revolutionized the treatment of cancer and antibodies targeting immune checkpoints or ligands thereof, *e.g.*, PD-1/PD-L1 or CTLA-4, have demonstrated clinical benefit. In particular, anti-PD-1/PD-L1 in head and neck or salivary gland tumors has shown promise (13–15). Through immune-expression of PD-1, PD-L1, major histocompatibility complex class I (MHC I), and the cancer testis antigen PRAME, a recent paper probed the TME of 53 SDCs (16). The investigators determined a correlation between the expression of PD-1, PD-L1, and PRAME. In addition, the expression of PD-1 in immune cells and PD-L1 in tumor cells were significantly associated with patient survival; and MHC I downregulation was observed in 82% of the SDCs. Taken together, these results indicate a potential for therapies targeted against the TME. Additional properties of the TME, *e.g.*, a desmoplastic stroma or immunosuppressive elements, may nonetheless limit the benefit of immunotherapies and should be investigated in detail.

In this study, the first comprehensive genomic characterization of SDC is reported. Transcriptomics and proteomics were exploited to unravel important molecular pathways that are involved in this cancer. In particular, several pathways related to extracellular matrix (ECM) remodeling, inflammation, and immunosuppression were apparent and have potential implications in personalized patient therapy. Based on genomic data, immunohistochemistry (IHC) and multiplexed immunofluorescence (IF), analysis of the TME composition enabled definition of two main SDC subtypes, revealed the importance of macrophages in SDC, and highlighted a number of opportunities to disrupt the SDC microenvironment.

## Materials and Methods

Additional details on the patient cohorts, proteomics, IHC and multiplexed IF are provided as Supplementary Material.

### Patients and cohorts

Published data for 16 SDC patients, referred to as the MSKCC cohort (see the original publication for details (5)), were combined with data from 20 additional patients from France and 4 from Belgium (Table 1). The patients diagnosed with SDC provided written informed consent for tissue collection and subsequent research. For the first cohort of 8 French patients (cohort 1), tissues from FFPE blocks plus snap frozen material obtained at the time of surgery were available. Normal salivary gland tissue from two patients were also collected. For the second cohort of 16 French and Belgian patients (cohort 2), only FFPE blocks were available. In addition, normal salivary gland tissue from three patients were provided; plus, for one patient, tissues from metastatic and primary SDCs were available.

**Table 1.** Clinical information of 24 SDC patients (cohorts 1 and 2).

| Clinical Feature | n (%) or mean (range) |
| --- | --- |
| Male | 16 (67%) |
| Age at diagnosis | 64 (33 - 87) |
| Cardiovascular risk | 11 (40%) |
| **Primary tumor site** |  |
| Parotid | 22 (92%) |
| Submandibular gland | 1 (4%) |
| Submaxillary gland | 1 (4%) |
| **T classification** |  |
| T2 | 8 (33%) |
| T3 | 4 (17%) |
| T4 (including 4a and 4b) | 12 (50%) |
| **N classification** |  |
| N0 | 7 (29%) |
| N1 | 2 (8%) |
| N2 | 1 (4%) |
| N2b | 11 (46%) |
| NX | 3 (13%) |
| **M classification** |  |
| M0 | 9 (38%) |
| M1 | 3 (12%) |
| MX | 12 (50%) |
| **Initial therapy** |  |
| Surgery+RT | 14 (58%) |
| Surgery+RT+CT | 8 (34%) |
| Surgery | 2 (8%) |
| **Disease Recurrence** |  |
| No | 6 (25%) |
| Local and regional | 1 (4%) |
| Local and distant | 3 (13%) |
| Regional | 1 (4%) |
| Distant | 8 (33%) |
| Deceased before recurrence | 2 (8%) |
| Recent cases (unevaluable) | 3 (13%) |
| **Second line therapy** |  |
| Radiotherapy | 1 (8%) |
| Chemotherapy | 5 (38%) |
| Surgery | 4 (31%) |
| Immunotherapy | 1 (8%) |
| Unknown | 2 (15%) |

### Transcriptomics

RNA was extracted from cohort 1 samples and quality-controlled. DNA libraries were prepared with the NEBNext Ultra II mRNA-Seq kit and sequenced on a HiSeq4000 (Illumina) using 2×75pb cycles to generate 130 million reads. MSKCC cohort transcriptomes (Fastq files) were downloaded from the NCBI Sequence Read Archive. With our pipeline, Fastq files of cohorts 1 and MSKCC were aligned against the human genome (Ensembl GRCh38) (STAR using default parameters and 2 passes, read counts extraction with HTSeq-count). See **Supplementary Material** and **Suppl. Table S1** for batch-effect correction and data normalization.

### Proteomics

Twenty FFPE tissue sections (5 µm thickness) were deparaffinized and the proteins digested by trypsin. Following digestion, 10% of each sample was mixed to create a pool for generation of the spectral library. The samples were analyzed on a nano-HPLC system (Sciex, Framingham, MA, USA) connected on-line to an electrospray Q-TOF 6600 mass spectrometer (Sciex). Two acquisition modes were used: data dependent (DDA) to generate the reference spectral library, and data independent (DIA or SWATH) to measure the samples. The library data was searched against the human protein database using Protein Pilot (Sciex). For the SWATH acquisition, the DDA method was adapted using the automated method generator embedded in the Analyst software (Sciex). Protein identification and quantitation was conducted with the Peak View software and the previously-generated protein library. Expression data for 3,203 proteins was obtained, and normalized against the total ion intensity before log_10_-transformation.

### Differential gene and protein analyses

Differential gene analysis was performed with edgeR (17), P-value < 0.01, FDR < 0.01, minimum fold-change 2, and minimum of 20 (normalized) read counts over all the samples analyzed. Differential protein expression was performed with limma (18), P-value < 0.05, minimum fold-change 1.25, and a minimum average (log_10_-transformed) signal of 3. Heat maps were generated with ComplexHeatmap (19). Dendrograms that ordered samples (columns) and genes/proteins (rows) were constructed with Ward’s method based on the Euclidean distance. For color assignment, a threshold of 2.5% was applied to the upper and lower values.

In conjunction with a hypergeometric test, GO terms and Reactome pathways were used. This was followed by Benjamini-Hochberg multiple hypothesis correction (internally-developed R script), FDR < 0.05 and a minimum of 5 query genes/proteins in a pathway or GO term.

### Immunohistochemistry

Paraffin sections (5 µm thick) were deparaffinized. Antigen retrieval was performed using AR6 buffer for 10 min in a pressure cooker. The sections were blocked for 30 min in protein block serum-free solution and incubated with the primary antibody at room temperature (RT) for 2h (see **Suppl. Table S2** for antibodies used). The slides were then washed and incubated for 30 min at RT with the secondary antibody. Subsequently, the sections were washed and then stained with 3,3’-diaminobenzidine (DAB). Stained sections were imaged with an automated Nanozoomer 2.0HT (Hamamatsu) at ×40 magnification (230 nm/pixel).

DAB-positive stained cells (CD3 and CD8 markers) were automatically counted using the open-source software Qupath (20). Due to particular cell shapes, macrophages (CD68, CD163) and α-SMA markers were evaluated by counting DAB positive pixels to improve accuracy. Intensity thresholds and other parameters for cell/pixel detection and classification were manually set for each staining type and performed identically for all samples. A machine learning-based method was applied to annotate stromal area and tumor nests within the tumor core. For further analyses, cell and pixel densities were estimated as the percentage of positive cells per mm² and the percentage of positive pixel per mm² of surface area, respectively (20). All steps were performed under the supervision of an expert pathologist (VC). Necrosis, tissue folds and entrapped normal structures were carefully removed.

### Multiplexed Immunofluorescence

Tissue sections were prepared as described above with the exception that incubation with the primary antibody that was conducted at 4°C overnight and the staining was performed using the Opal system (Perkin Elmer). Following primary antibody incubation, the slides were incubated with the corresponding secondary antibody as described above. The slides were then incubated with 100 μL staining solution prepared from 2 μL Opal dye and 98 μL amplifying buffer. Following 10 min incubation, the slides were washed and subjected to microwave-assisted antibody removal. After cooling and a wash in PBS buffer for 5 min, the tissues were re-blocked. Tissues were then incubated with the next primary antibody and the staining procedure was repeated using the following Opal dyes: 520, 570, 620 and 690.

Multiplexed immunofluorescence (IF) images were treated with Fiji software and analyzed with an internally-developed R script. The images were converted to 8-bit gray scale; a lower threshold was applied to remove the background noise and an upper threshold was applied to rescale the maximum gray value. The IF image of the receptor was binarized to isolate the cells expressing the receptor. The *Analyze Particles* plugin of Fiji was used to locate the cells and extract the positions of the centroid. Only the shapes with a circularity > 0.3 and an area > 50 pixel² were retained. The R script used these positions to calculate the average fluorescence of the receptor inside a circle with diameter equal to a receptor-expressing cell and the average fluorescence of the ligand inside a surrounding crown (**Suppl. Fig. S1A**). The averaged fluorescence values were then used to calculate an IF ligand-receptor score (ifLR-score). A threshold on the ifLR-score was determined to assess if the interaction was positive for each cell expressing the receptor (**Suppl. Fig. S1B**). See Supplementary Materials for details of the score calculation and threshold adjustment.

### Ligand-receptor interactions and pathways

Interactions from FANTOM5, HPRD, HPMR, the IUPHAR/BPS guide to pharmacology, UniprotKB/Swissprot annotations, Reactome, plus manual extraction from cellsignaling.com maps and the literature were combined. Reactome-derived ligand-receptor (LR) pairs corresponded to protein interactions from Reactome with the respective participants annotated as ligand or receptor in Gene Ontology (GO). Reactome pathways that were downloaded as a collection of binary interactions from PathwayCommons, were also used. For a few receptors that remained unconnected in Reactome, these were manually complemented with interactions annotated in UniprotKB/Swissprot (**Suppl. Table 3**).

Confident LR interactions that occur in SDC were determined by firstly imposing Spearman r > 0.5 between a ligand and its receptor and Benjamini-Hochberg corrected P-values of these correlations < 0.01. This resulted in 277 LR pairs. Receptor downstream activity was then assessed by considering all Reactome pathways containing the receptor. In each pathway, the target genes were identified plus otherwise controlled genes, *e.g.*, by phosphorylation of the product, and the criterion that at least 4 displaying Spearman r > 0.5 with the receptor was imposed. If only 2 or 3 targets/controlled genes were available, r > 0.5 was required for all. This procedure selected 151 confident LR pairs. The same algorithm was applied for GO biological process (GOBP) terms as *ad hoc* pathways. Their topology was retrieved from Reactome interactions and 144 confidence LR pairs were obtained. In total, 179 unique confidence LR pairs were determined.

## Results

### Functional genomics of salivary duct carcinoma

The availability of non-tumor tissues in our two cohorts enabled us to ask the fundamental question: which are the pathways that are deregulated in SDC and whether any of these can be targeted. This investigation was initiated at the transcriptional level by combining the RNA-seq data from the MSKCC cohort and our cohort 1. The merged data sets were submitted to the same bioinformatic pipeline and non-tumor tissues *versus* SDCs were compared. This revealed 1,634 genes that were significantly-regulated (Fig. 1A); the majority (1,451) with increased expression. Similarly, the second cohort was exploited to perform a similar comparison at the protein level. This resulted in 244 significantly-regulated proteins, 213 with increased expression (**Suppl. Fig. 2**). Pathway enrichment analysis was conducted separately on the regulated genes and proteins. Beyond pathways commonly regulated in tumors (transcription, cell cycle, *etc.*), a number of more SDC-relevant pathways were selected. These are featured in Fig. 1B (complete tables are provided as **Supplementary Data**).

**Figure 1.**
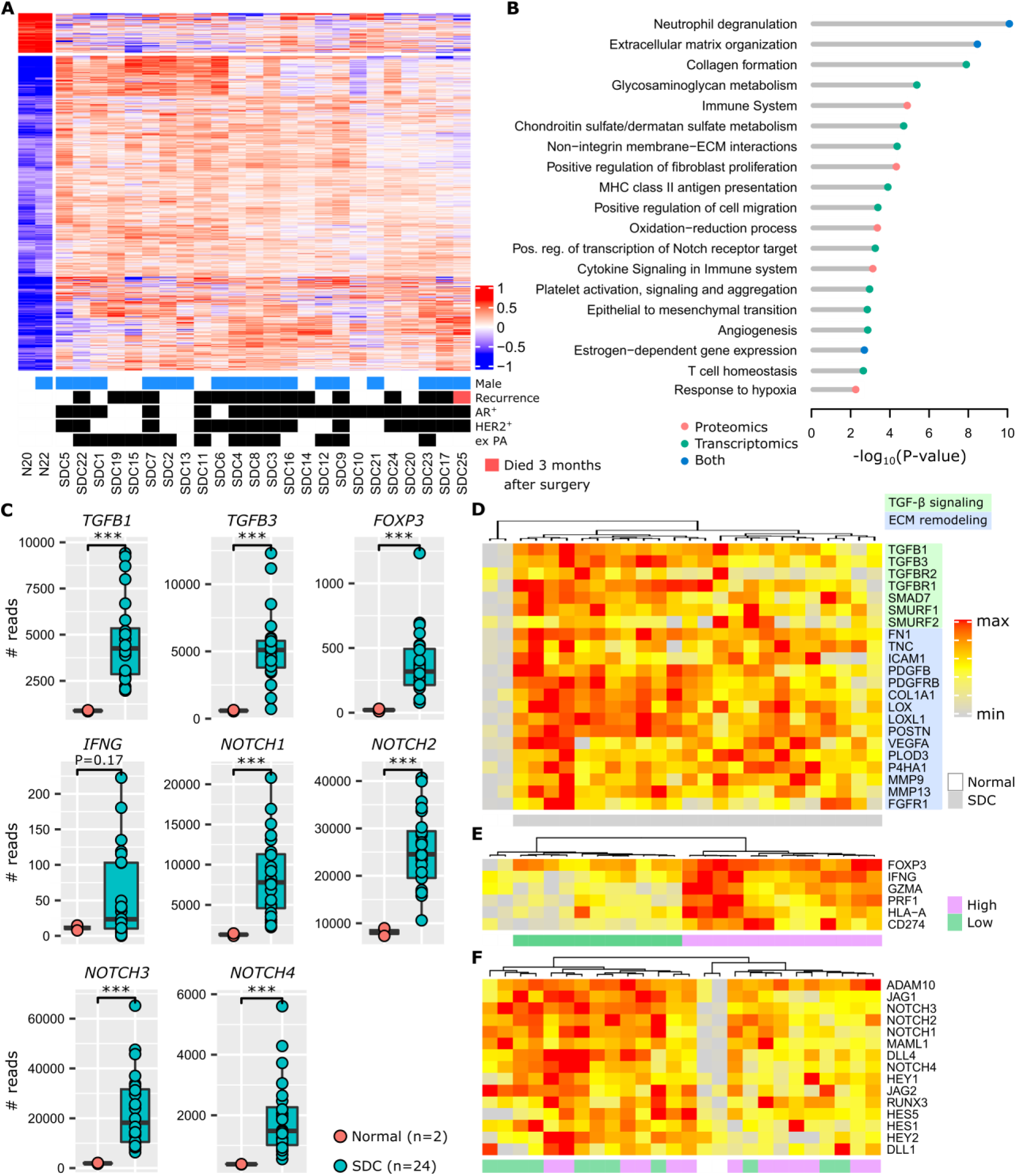
Functional genomics. **A**. Differentially-expressed genes that compare adjacent normal tissues with SDC. **B**. Selected pathways. **C**. Expression of important genes (Wilcoxon one-sided test, n=26=2+24, *** P<0.005). **D**. TGF-β and ECM remodeling genes. **E**. Inflammation and immunosuppression genes. Two clusters of tumors exist, denoted high and low. Expression of *FOXP3* is high in all SDCs. **F**. Notch signaling genes. Note that the gradient (from left to right) of Notch signaling gene expression is not correlated with clusters in panel F.

The invasive component of SDC presents as a desmoplastic stromal reaction (DSR) with a partially-hyalinized ECM. This is consistent with ECM remodeling, collagen formation, glycosaminoglycan and chondroitin metabolism, integrin interactions, and fibroblast proliferation pathways that were increased. ECM remodeling contributes to fibrosis, tumor stiffness, and to a permissive environment for neo-angiogenesis plus tumor cell spreading. Several additional regulated pathways indicated that these processes occur in SDC: epithelial to mesenchymal transition (EMT), angiogenesis, and hypoxia. Of particular interest, TGF-β expression was strongly augmented (Fig. 1C), and is known to contribute to an immunosuppressive TME (21,22). In Fig. 1D, the activation of TGF-β signaling is illustrated; plus the activity of several genes involved in ECM remodeling. Depending on the SDC, but compared to normal tissues, different degrees of expression for these genes was observed. Overall, however, all were clearly overexpressed.

Fig. 1B shows an increased expression of pathways related to an inflammatory response (neutrophil degranulation, MHC class II antigen presentation, cytokine signaling) together with immunosuppression (T cell homeostasis). This is supported by *FOXP3* upregulation (Fig. 1C), a transcription factor expressed by regulatory T cells (Tregs). An increase in interferon-γ (*IFNG*), a cytokine that can activate macrophages and NK cells, is also apparent. Interferon-γ adopts a pro-tumoral and immunosuppressive role in certain tumors (23) and, within the limits of the small cohort 1, a significant association of *IFNG* abundance with relapse was indeed observed (**Suppl. Fig. S3**). Representative genes for inflammation and immunosuppression are featured in Fig. 1E, *e.g.*, T cell effectors such as perforin (*PRF1*) and granzyme A (*GZMA*) or the immune checkpoint ligand PD-L1 (*CD274*). Two groups of SDC are suggested by the gene expression pattern: high *versus* low level of inflammation/immunosuppression. For each SDC, *FOXP3* ubiquitous expression is consistent with TGF-β signaling.

Notch family members appear to be important in SDC (Figs. 1C), and many downstream genes of the Notch signaling pathway were overexpressed (Fig. 1F). Notch contributes to cancer cell development but also the dialog and interaction with the TME (24). This is similar to ECM-remodeling and TGF-β signaling, however, a gradient of Notch signaling intensity can be seen across the SDCs in Fig. 1F. The activity of the pathway was strongly augmented in every tumor.

Activation of estrogen-dependent gene expression correlates with the apocrine-like nature of SDC transcriptomes (5) and the frequent AR+ histological status (~80%). Contrary to breast cancer, clear and coherent transcriptional subtypes were not observed (Fig. 1A and **Suppl. Fig. 2**). Correlation of protein and gene expression with clinical data (recurrence, HER2 or AR-positive staining, ex PA origin) was attempted, however, no significant or biologically-sound association was observed.

### Probing the microenvironment of salivary duct carcinomas

A software tool used to score cell populations of the TME from transcriptomic data, MCP-counter (25), was applied. Due to the modest size of our cohort, and the histological proximity of SDC with breast IDC (1), 624 breast IDCs were retrieved from the cancer genome atlas (TCGA) and were submitted to MCP-counter together with the SDCs. With respect to a large number of tumors, this enabled normalization of the MCP-counter score for each TME cell type. Depending on the number of immune cells, SDC clearly formed two distinct clusters (Fig. 2A). Normalized fibroblast and endothelial cell abundances are represented, but these neither contribute to the dendrogram nor correlate with the clusters. Cluster 1 (n=12) was enriched for immune cells: T and B lymphoid cells and monocytic lineage cells. Cluster 2 (n=14) was represented by low lymphoid cell infiltrate but contained heterogeneous amounts of cells from the monocyte lineage. Due to clinical relevance, MCP-counter estimates were validated by IHC (using CD3 and CD8 antibodies) on the 8 SDCs from cohort 1 (Figs 2B). Differential gene expression between cluster 1 and 2 confirmed a strong and almost exclusive variation in immune pathways (**Suppl. Fig. S4**). Hence, cluster 1 was termed *immune-infiltrated* and cluster 2, *immune-poor*. The 8 SDCs from cohort 1 together with 14 SDCs from cohort 2, for which we also performed IHC, were classified with respect to CD3- and CD8-positive cell enumeration and localization (Fig. 2C and **Suppl. Fig. 5**). In 8/22 cases (36%), a very low number of T cells were observed (Fig. 2D); these comprised the immune-poor group. Two patterns of immune-infiltrated SDC were apparent (Fig. 2D). In 4/22 (18%) tumors, CD8+ T cells were restricted to the invasive margin (IM). These have been previously-characterized as T cell-excluded (26), and are defined here as the *immune-infiltrated IM* group. Another group of tumors, 10/22 (46%), displayed T cells either in the tumor core (TC) only or, in the TC and at the IM. Referred to as the *immune-infiltrated TC* group, a quasi-exclusion of CD8+ T cells from the tumor nests in these 10 cases was observed. This particular pattern has been previously-described in lung, pancreatic, and ovarian carcinomas and was also characterized as T cell-excluded (27).

**Figure 2.**
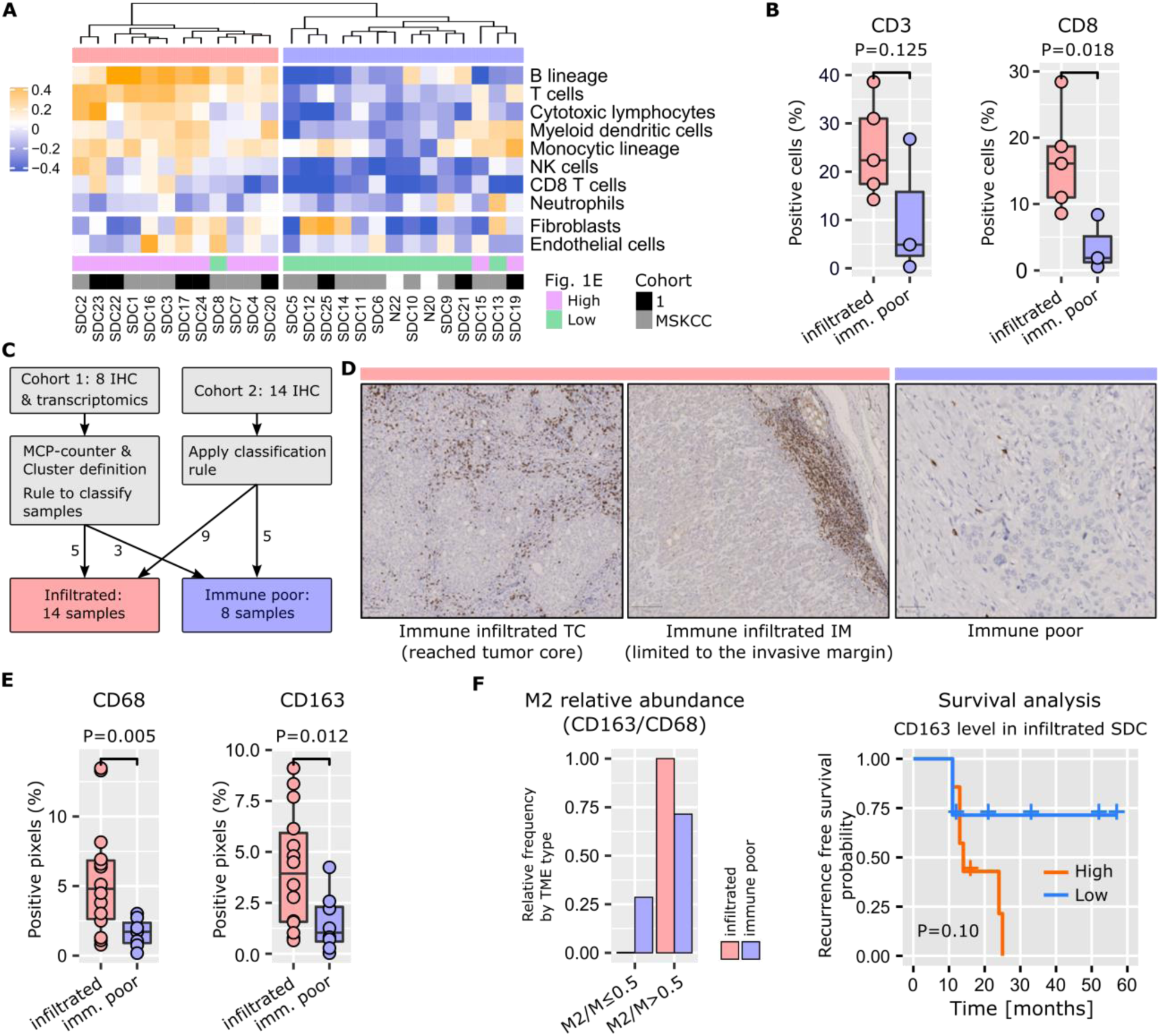
TME cell types. **A**. Application of MCP-counter to SDC transcriptomes revealed two groups of SDC: immune-infiltrated (light red) and immune-poor (light blue). Note that these two groups are almost identical to the high/low clusters in Fig. 1E. **B**. Validation of the differential T cell infiltrates between the groups (Wilcoxon one-sided tests, n=8=5+3). **C**. Samples from cohort 2 that were not available for transcriptomics were added to the IHC study to obtain 22 SDCs. Samples were classified according to CD3+ and CD8+ cell abundance and localization. **D**. Two CD3+/CD8+ patterns were observed with the immune-infiltrated group: limited to the IM, or present in the TC. **E**. Total macrophages (CD68) and M2 (CD163) were more abundant in the immune-infiltrated SDC (specific pattern ignored), (Wilcoxon one-sided test, n=21=14+7 for CD68, one immune-poor outlier removed (significant for Grubbs and Dixon tests, robust (median and MAD) z-score>3); n=22=14+8 for CD163). **F**. Distribution of SDC with M2 macrophages representing > 50% of the macrophages.

Tumor-associated macrophages (TAMs) play an important role in TME homeostasis and resistance to various treatments (26,28–30). MCP-counter indicated a variable presence of TAMs in the two SDC groups (monocytic lineage in Fig. 2A); therefore, TAM density within the stromal areas of the TC was assessed by IHC and digital imaging. Macrophages and the alternative activated phenotype (called M2) were stained for CD68 and CD163, respectively. A significantly-higher TAM content in the immune-infiltrated SDC was observed (Fig. 2E). The proportion of stromal M2 macrophages (M2/M ratio) was based on the CD163 density/CD68 density ratio. A high M2/M ratio (> 0.5) was seen in 6/8 (75%) of the SDCs with an immune-poor phenotype and 13/13 (100%) with the immune-infiltrated phenotype. This suggested that M2 macrophages represent a significant proportion of TAMs in SDC (Fig. 2F). With respect to M2 abundance in the TME, recurrence-free survival (RFS) analysis indicated a clear trend. Previous results in breast, pancreatic, and oral cancers are in agreement with this finding (29–31); thus indicating potential relevance of TAM-targeting therapies for SDC.

As previously mentioned, SDC histology is characterized by a dense stroma. The DSR was assessed by measuring α-SMA stromal density and graded as follows: grade 0 (< 5%), 1 (5-15%), 2, (15-50%), 3 (> 50%). The data revealed that 22/29 (86%) SDCs displayed a DSR grade ≥ 2 (Fig. 3). Interestingly, the DSR was independent of the immune infiltrate (**Suppl. Fig. 6**) and; in our data, there was a trend between α-SMA levels and RFS (Fig. 3).

**Figure 3.**
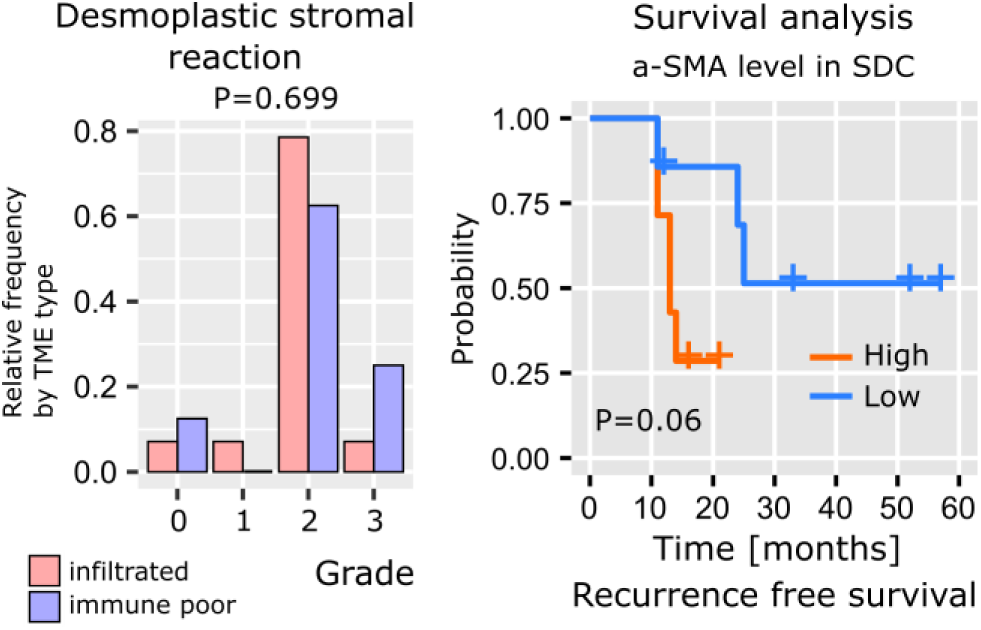
SDC desmoplastic stromal reaction. The distribution of DSR grades is comparable between immune-infiltrated and immune-poor SDC (Kolmogorov-Smirnov test, n=22) and recurrence-free survival (Kaplan-Meier curve, log-rank test, n=22, high=above median, low=below median).

### Mapping cellular interactions in the SDC stroma

The cellular network of a tumor is significantly rewired compared to the original healthy tissue. A typical illustration is the induction of PD-1 positive T cell inhibition and functional exhaustion thereof by PD-L1-expressing epithelial cancerous cells and/or immune infiltrated cells (32). It is reasonable to suggest that a systematic study of cellular interactions within the SDC TME may unravel elements that can be potentially targeted. Ligand-receptor (LR) interactions from several public databases and the literature were compiled to assemble a database (LR*db*) comprised of 3,270 unique LR pairs. Next, an algorithm (Fig. 4A) was developed to search for evidence of these interactions in SDC transcriptomes. Briefly, each LR pair in LR*db* was assessed and a Spearman correlation r > 0.5 was imposed between a ligand and a receptor to result in 277 filtered LR pairs. Evidence for receptor downstream activity was then assessed using Reactome pathways and additional correlations. This procedure (Materials and Methods) selected 179 confident LR pairs (**Supplementary Data**). Our algorithm associates each receptor with pathways, the recurrent pathways summarized in Fig. 4B largely mirror the TME-associated pathways in Fig. 1B.

**Figure 4.**
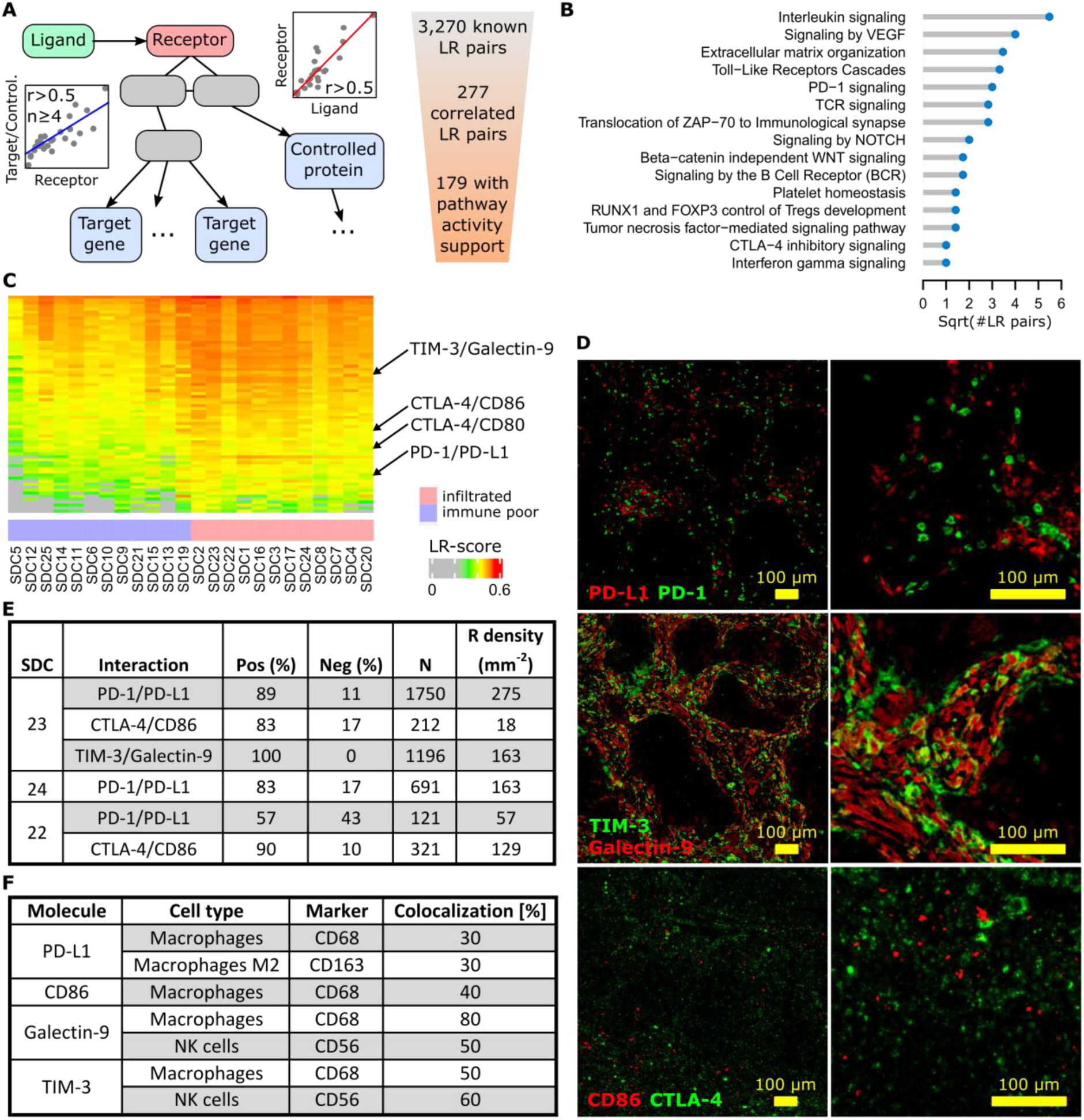
Ligand-receptor interactions within the SDC microenvironment. **A**. Principle of the LR pair-search algorithm: the ligand and the receptor are required to correlate over the SDC transcriptomes, and the candidate receptor has to occur in pathways with at least 4 downstream regulated genes (transcription factor targets) or regulated proteins (phosphorylation or other PTM) that display sufficient correlation with the latter receptor. **B**. Functional categories where the ligands and the receptors occur. **C**. Ligand-receptor pairs with transcriptional LR-score correlated with the immune-infiltrate category (72 pairs). **D**. The three selected pairs in SDC23. **E**. Ligand-receptor co-localization results. **F**. Semi-quantitative assessment of the degree of fluorescence for the ligand (PD-L1, galectin-9, CD86) or receptor (TIM-3) colocalized with the marker fluorescence for each assessed cell type (note that for galectin-9, the fluorescence of CD56+ and CD68+ cells sums to 130%, but these measurements were obtained from different slides and are semi-quantitative only).

### Targets for immunotherapy

The search for immune infiltrate-related LR pairs was achieved by computing a score (the LR-score) that reflected the co-occurrence of the ligand and the receptor in each SDC transcriptome (Materials and Methods). A Spearman r > 0.6 was imposed with MCP-counter immune cell gene signatures to select 72 LR pairs (Fig. 4C and **Suppl. Fig. S7** with LR pair names). From these pairs, three immune checkpoints were chosen for validation by multiplexed IF: PD-1/PD-L1, CTLA-4/CD86, and TIM-3/galectin-9 (Fig. 4D). By IF, the percentage of cells expressing the receptor that were in proximity to cells expressing the ligand were counted. Thus, the signal from the extracellular ligand overlapped with the receptor signal at the membrane. This enabled the determination of average cell diameters, the definition of a crown-shaped signal overlap area, and the computation of a score in this area; *i.e.*, an analog of the LR-score (Materials and Methods).

#### PD-1/PD-L1 interaction

In three SDCs classified with the immune-infiltrated phenotype, a potential interaction between PD-1 and PD-L1+ cells was investigated. Two were from the TC group (SDC23 and SD24) and one was from the IM group (SDC22). For SDC23 and SDC24, a large percentage of PD-1+ cells adjacent to PD-L1+ cells were observed; 89% and 83%, respectively (Fig. 4E). In SDC22, this percentage (57%) was lower although superior to 50%. The results indicated that a substantial proportion of PD-1+ cells were exhausted in the TME of immune-infiltrated SDCs. This may explain the absence of an association between CD8+ T cells and recurrence-free survival with such tumors (P=0.94, see **Suppl. Fig. S8**). It was also noted that SDC22 (immune-infiltrated IM phenotype) contained less PD-1+ cells. The expression of PD-L1 by M2 (CD163) and non-M2 macrophages (Fig. 4F) was further shown. A proportion of the fluorescence could not be attributed to macrophages and was assumed to originate from other cells.

#### CTLA-4/CD86 interaction

CTLA-4 is an immune checkpoint, and the expression thereof in SDC has never been studied. Inhibition of CTLA-4+ cells by adjacent CD86+ cells (TAMs) was assessed in SDC23 and 22. In both cases, the percentage of positive instances was high; 83% and 90%, respectively. The density of CTLA-4+ cells, however, was less than PD-1+ cells. This was particularly evident with SDC23 (Fig. 4E). CD86 was expressed by CD68+ cells but not only by this cell type (incomplete attribution of IF).

#### TIM-3/Galectin-9 interaction

T-cell immunoglobulin mucin receptor 3 (TIM-3, *HAVCR2* gene) is an immune checkpoint that plays a role in T-cell exhaustion, and binding to galectin-9 (*LGALS9* gene) suppresses T cell response. The TIM-3/galectin-9 interaction was assessed in SDC23 and 100% of cells were observed to express TIM-3 adjacent to galectin-9+ cells (Fig. 4E). Galectin-9 co-localized with CD68+ (macrophages) and CD56+ (NK) cells (Fig. 4F). As mentioned above, these two cell types do not explain the total signal from galectin-9; thus indicating that other cells also express the ligand. Strikingly, co-localization of TIM-3 with CD68 and CD56 gave the same results. The suggestion from this is that there is potential (paracrine) cross-inhibition of macrophages and NK in addition to autocrine inhibition. IF images contained clear examples of each cell type expressing both galectin-9 and TIM-3 (**Suppl. Fig. S9**). From the literature, it is known that TIM-3 overexpression is observed in NK and macrophages in advanced tumors (33,34).

## Discussion

A functional genomic survey was conducted where SDC was compared to adjacent non-tumor tissue by proteomics and transcriptomics (Fig. 1). This analysis revealed numerous genes involved in ECM remodeling and CAF proliferation that not only facilitates progression towards aggressiveness, EMT, and angiogenesis; but also induces the simultaneous expression of inflammatory and immunosuppressive pathways. Notch and TGF-β signaling were activated and probably contribute to such a TME (21,22,24). In mice, overexpression of TGF-β in normal salivary glands indeed causes ECM remodeling and the replacement of normal glandular parenchyma with interstitial fibrous tissue (35).

By combining immune cell gene signatures and T cell IHC quantitation and localization (CD3 and CD8), two groups of SDC were defined: immune-poor (36% of SDCs in our study) *versus* immune-infiltrated (64%) (Fig. 2). An increase in TAM concentration was observed in the infiltrated group, where more than 50% of the macrophages were M2. As observed with other tumors (36), a clear trend was shown between M2 abundance and RFS in immune-infiltrated SDC (Fig. 2F). Depending on the localization of the T cells, *i.e.*, present in the TC (46%) or restricted at the IM (18%); two sub-groups for immune-infiltrated SDC exists. When T cells were present in the TC, these were concentrated and maintained at the periphery of tumor nests, a structure that has been previously-observed and could be due to TAMs and stromal cells (27,28). Based on T cells, the immune poor group resembles immune-deserted tumors (26,37). The characterization of SDC immune-infiltrate phenotypes establishes a first concept for patient segregation with respect to immunotherapy. Immune-poor tumors are obviously less likely to benefit from such treatments and the importance of TAMs in infiltrated SDC should not be disregarded.

Given the strong ECM remodeling signature determined by transcriptomics and proteomics, SDC DSR was assessed by measuring α-SMA abundance. As expected, a large proportion of SDC with DSR grades ≥ 2 (86%) was apparent. Interestingly, this was independent of the immune-infiltrate phenotype (**Suppl. Fig. S6**). Moreover, across all the SDCs, there was a clear trend between α-SMA abundance and RFS (Fig. 3). These results clearly show that the physical barrier opposed by SDC-dense stroma should be addressed for both the infiltrated- and poor-immune phenotypes.

Opportunities of therapeutic disruption of SDC TME can be surveyed through the generation of a map of cellular interactions that can be targeted. Hence, an inference algorithm was developed that identified 179 confident predictions of active LR pairs that covered immune and non-immune functions (Fig. 4). From this list, 72 LR pairs were identified that correlated with SDC-immune phenotypes that depicted a complex network of mixed pro- and anti-inflammatory interactions (**Suppl. Fig S7**). Three immune checkpoints with existing inhibitory molecules were selected for validation by multiplexed IF. These were: PD-1/PD-L1, that has already been evaluated in SDCs (16); CTLA-4/DC86; and TIM-3/galectin-9. The latter two have never been previously evaluated for SDC (Fig. 4D). The three interactions were confirmed (Fig. 4E) and it was shown that each ligand is produced by TAMs (Fig. 4F). Thus, further evidence was provided that this immune cell type plays an important role in SDC immunosuppression. For the TIM-3/galectin-9 interaction, NK cells were additionally identified as a source of ligand that is consistent with interferon-γ expression (Fig. 1). In fact, the data even suggested a complex self- and cross-inhibition of macrophages and NK cells (**Suppl. Fig. S9**) that, upon disruption, could unleash concomitant antitumor reactions. Indeed, monoclonal antibodies against galectin-9 reduced TAMs towards an M1 phenotype *in vitro* (38); whereas TIM-3 blockade increased intratumoral NK cell cytotoxicity in mice with MHC class I-deficient tumors (34) similar to SDCs (16).

Our results have obvious clinical implications. The definition of two major immune phenotypes suggests that patients could be stratified after initial surgery, as is the practice for many tumors (26,37,39). In agreement with the absence of CD8+ cell abundance and RFS, the validated PD-1/PD-L1 interaction indicates exhaustion of effector T cells (37) (**Suppl. Fig. S8**). Furthermore, the many additional immunosuppressive LR interactions predicted in our study, including validated TIM-3/galectin-9 and CTLA-4/CD86, suggest that single immunotherapies are unlikely to satisfactorily treat SDC tumors. Rather, combined approaches should be favored, *e.g.*, anti-PD-1/PD-L1 and anti-CLTA-4. Given the role of NK cells, strategies exploiting the innate immune system should also be considered (34,40). In addition, the dense stroma of SDC cannot be ignored and could be addressed by the proposed regimen. For instance, in a murine pancreatic ductal adenocarcinoma (PDAC) model, depletion of FAP-expressing CAFs synergized with anti-PD-L1 immunotherapy (41). In metastatic urothelial cancers, blocking TGF-β also resulted in an improved efficacy of anti-PD-L1 antibodies by reducing TGF-β signaling in stromal cells and improving the penetration of T cells in the tumor (21). In PDAC, it has also been proposed that focal adhesion kinase inhibitors can be used to improve checkpoint immunotherapy efficacy (42). The interest of addressing the stroma could be even higher for SDC devoid of immune infiltrate. Targeting hyaluronic acid (HA) by PEGPH20, a pegylated recombinant human hyaluronidase, resulted in an improved delivery of small molecule therapy in PDAC (43). Phase 2 and 3 trials of PEGPH20 plus chemotherapy are underway in metastatic PDAC. Our functional genomic data indicated that combinations with Notch or drugs against specific mutations (5) could be successful.

The validation of LR pairs by multiplexed IF is tedious work. Nevertheless, successful validation of the three selected pairs encouraged us to open this discussion to additional inferred LR pairs. In particular, pairs where literature exists for other tumors would be worth pursuing for SDC. As an example, the leukocyte immunoglobulin-like receptors *LILRB1* and *LILRB2* have been observed to interact with HLA class I molecules and correlate with immune and monocytic lineage gene signatures in transcriptomics. Up-regulation of either LILRB1 or LILRB2 in macrophages was shown to provide an evasion mechanism for cancer cells against phagocytosis. Through activation of AKT and IL-4 signaling, LILRB2 antagonism induced a reduction in PD-L1 expression by macrophages and reprogramming of lung TAMs (44). CCR5 and ligands thereof (CCL3, CCL4, CCL5, CCL8, CCL11, and CCL13) were observed to follow the same pattern with a higher LR-score in immune and monocytic lineage infiltrated SDCs. CCR5 interactions were reported to potentiate the recruitment of CCR5-expressing TAMs plus the immunosuppressive characteristics. Together with enhanced DNA repair, CCR5 signaling may induce a pro-inflammatory and pro-metastatic immune phenotype and thereby conveying resistance to DNA-damaging agents (45). CCR5 inhibitors (maraviroc and leronlimab) were approved by the FDA, and in combination with either immune-checkpoint inhibitors or chemotherapy for colorectal cancer (NCT01736813, NCT03274804, NCT03631407) and TNBC (NCT03838367), clinical trials are currently underway. OX40L (TNFSF4) and the receptor (TNFRSF4) expression were correlated with fibroblasts, T cells and the monocytic lineage. In triple-negative breast cancer (TNBC), OX40L was recently shown to be expressed by a subgroup of fibroblasts (α-SMA, PDGFRB, FAP, CAV1 positive myofibroblasts) that can attract Tregs in the TME. By subsequent interaction, this results in an enhanced capability to inhibit the proliferation of effector T cells (46). Consequently, OX40 agonists should not be used in SDC as these may further impair the immune response. The ensemble of our inferred, confident LR pairs is provided as **Supplementary Data**, including references to known drugs and current clinical trials.

This study is the first attempt to profile the SDC microenvironment and combine the information gained with functional genomics via a comprehensive exploitation of transcriptomics, proteomics and digital imaging. It proposes a large repertoire of cellular interactions that could be disrupted, and thus complements existing research by others who characterized the actionable mutations of this tumor. Based on immune infiltrate, two groups of SDC were identified; and this should provide a rationale for patient enrolment in clinical trials. The importance of macrophages, NK cells and potentially Tregs, was also shown, and should also be taken into consideration to better define patient groups. Moreover, upregulated pathways that could be targeted, *i.e.*, Notch and TGF-β, were discovered. The clear trends between M2 macrophage or α-SMA abundance and DFS could be developed into tools to manage patients following tumor resection. In addition, our work unraveled multiple novel options for SDC devoid of immune infiltrate by targeting the stroma.

## Acknowledgements

We thank the Réseau d’Expertise Français des Cancers ORL Rares (REFCOR) and the Centre Ressources Biologiques (CRB) of Montpellier University Hospital (CHU) for sample access. We also thank Prof. Olivier Adotevi (UMR 1098 Inserm, Besançon) for very valuable input during the development of this project and Prof. Luc GT Morris (MSKCC) for assisting us in using his data.

