## Supplemental Information for "The molecular landscape and microenvironment of salivary duct carcinoma reveal new therapeutic opportunities"

[1] Institut de Recherche en Cancérologie de Montpellier (IRCM), INSERM, Parc Euromédecine, 208 rue des Apothicaires, 34298 Montpellier, France
[2] Biological Hematology Department, CHU Montpellier, Hôpital Saint Eloi, 34275 Montpellier, France
[3] Université de Montpellier, Faculté de Pharmacie, 15 avenue Charles Flahault, 34093 Montpellier, France
[4] Institut Régional du Cancer Montpellier (ICM), Parc Euromédecine, 208 rue des Apothicaires, 34298 Montpellier, France
[5] Université de Montpellier, Faculté de Médecine, 2 rue école de Médecine, 34060 Montpellier, France
[6] Biopathology Department, CHU Montpellier, Hôpital Gui De Chauliac, 34000 Montpellier, France
[7] Biothèque, Université de Liège, 4000 Liège, Belgium
[8] Pathology Department, CHU Liège, Université de Liège, 4000 Liège, Belgium
[9] Université de Montpellier, 163 rue Auguste Broussonnet, 34090 Montpellier, France
[10] Department of Molecular Pharmacology and Oncology, Gunma University Graduate School of Medicine, Gunma, Japan
[11] Equal contribution

**Running title:** Molecular landscape and microenvironment of salivary duct carcinoma

**Keywords:** salivary duct carcinoma, stroma, personalized medicine, immunotherapy, molecular pathways

### Supplementary Materials and Methods

#### Patient consents

The French patients with SDC diagnosis were consented for tissue collection and research analysis under institutional reviewing board approval at the University Hospital of Montpellier (France). For the Belgian patients, the ethical committee of the University Hospital Liege has approved the use of human material in the current study. All samples were obtained from the institutional biobank of the University Hospital Liege, Belgium. According to Belgian law, patients obtained the information that the residual material could be used for research purpose and the consent is presumed as long as the patient does not oppose (opting-out), which was not the case for those patients.

#### Transcriptomics

Cohort 1 fresh frozen sample RNA was extracted using DNeasy Blood and Tissue Kit (Qiagen) and quantified by spectrophotometry using the Nanodrop 2000 (ThermoFisher Scientific). RNA quality and integrity was analyzed by 2100 Bioanalyzer (Agilent) and Fragment Analyzer. DNA libraries were prepared with the NEBNext Ultra II mRNA-Seq kit. Quantification of the library was obtained by real time PCR. Sequencing and data processing methods are detailed in the main paper.

PolyA+ selection and rRNA depletion are the main approaches for RNA preparation in RNA-seq studies. We used polyA+ selection and the MSKCC cohort was prepared with RiboErase rRNA depletion. Among coding genes, histone genes are known to be difficult to quantify with polyA+ selection whereas the other genes are quantified almost identically with the two approaches (1). In fact, searching for 50-fold or higher variation between cohort 1 and cohort MSKCC after upper quartile read count normalization yielded a list of 41 genes that was comprised of histones mostly:

**Supplementary Table S1. Most affected genes by differences in sample preparation, cohorts 1 *vs.* MKSCC.**

| | | | |
|---|---|---|---|
| AC009022.1 | HIST1H2AI | HIST1H3C | HIST2H2AA4 |
| AC087392.1 | HIST1H2AJ | HIST1H3F | HIST2H2AB |
| AL138751.1 | HIST1H2AL | HIST1H3I | HIST2H2AC |
| AL139333.1 | HIST1H2BB | HIST1H3J | HIST2H3A |
| CTD-2116N17.1 | HIST1H2BI | HIST1H4A | HIST2H3C |
| HIST1H1A | HIST1H2BL | HIST1H4B | HIST2H4B |
| HIST1H1B | HIST1H2BM | HIST1H4C | OR2D2 |
| HIST1H1D | HIST1H2BO | HIST1H4D | OR6A2 |
| HIST1H1E | HIST1H3A | HIST1H4F | TAS2R50 |
| HIST1H2AB | HIST1H3B | HIST1H4L | UBQLN3 |
| HIST1H2AH | | | |

We decided to remove these 41 genes from the study. The final read count matrix combining cohorts 1 and MSKCC was filtered by only keeping the genes expressed with > 5 reads in > 5 samples (16,680 genes). Subsequently, the matrix was normalized by total read counts. We observed no significant batch effect between cohorts in all our analyses and hence decided not to apply further data transformation.

**Proteomic Analysis of FFPE Samples**

Twenty FFPE tissue sections of five-micrometer thickness were deparaffinized with 1ml of xylen at 60°C for 10 min. Following this, the samples were centrifuged at 20.000g, room temperature (RT) for 5 min and the supernatant was removed. The xylen treatment was reapplied for a total of 3 times. Next, 1ml of ethanol was added and the samples were vortexed and centrifuged at 20.000g, RT for 5 min. The supernatant was discarded and the ethanol wash was re-applied to the pellet for a total of 4 times. The samples were then dried using Speed Vacuum and suspended in 500µl of citrate buffer (pH 6) with 1% SDS. Following a sonication step, the samples were incubated for 30min at 95°C under vigorous shaking. Next, the samples were allowed to cool down at RT for 20-30min and the pH was re-adjusted to 8.5 with 100mM NaOH solution. The samples were centrifuged (20.000g, RT for 5 min) and the supernatant was transferred to a new tube. The protein content of the resulting solution was determined using BCA Protein Quantification Kit (Thermo Fisher, Waltham, MA, USA; cat. no.: 23225). Hundred microgram of protein extract was transferred in a fresh tube and subjected to reduction using 20mM DTT for 30min at 60°C. Following this, the protein samples were alkylated using 50mM 2-chloroacetamide for 30min at RT, in the absence of light and under shaking. The proteins were then precipitated using 2D Clean-Up kit according to the manufacturer's instructions (GE Healthcare, Chicago, IL, USA; cat. no. 80648451). The protein pellets were then suspended in 50µl of ammonium bicarbonate 100mM /calcium chloride 1mM buffer (pH 8). To this suspension, 0.01% of Protease Max surfactant was added along with 1µg of trypsin. The samples were digested overnight (ON) at 37°C. Following digestion, 10% of each sample was transferred in a new tube where all the samples were mixed in a library. The remainder of the sample was Speed Vac to dryness. The library sample was further subjected to peptide fractionation using High pH Reversed-Phase Peptide Fractionation Kit according to the manufacturer's instructions (Thermo Fisher; cat. no.: 84868). From the library sample, 8 individual peptide fractions were derived, which were then Speed Vac to dryness. All the samples, including the library samples, were dissolved in 0.1% TFA and were subjected to salt removal using ZipTip according to the manufacturer's instructions (Merck, Darmstadt, Germany; cat. no.: C5737).

The peptide samples were analyzed using a 1D-nano-HPLC system (Sciex, Framingham, MA, USA), which was connected on-line with an electro spray Q-TOF mass spectrometer 6600 (Sciex). A total of 1 µg of sample was injected on the C18 analytical column (Acclaim® 75 µm x 150 mm, p/n: 162224; Dionex, California, USA) with a gradient of 0–40% phase B (90 % acetonitrile, 9.9 % water and 0.1 % formic acid) for 100 min at the flow rate of 0.3 µl/min. Two acquisition modes were used, data dependent (DDA) for the measurement of the library and data independent (DIA or SWATH) for the samples. In the DDA mode, mass spectral data were acquired over a mass range from 400 to 1600 m/z. One full MS scan was automatically followed by up-to 30 MS/MS scans of the most intensive peptides found in this mass range (bearing +2 or +3 charges). The acquired data for each fraction of the library sample were merged and used for MS/MS database search with Protein Pilot software (Sciex). For the SWATH acquisitions, the DDA method was adapted using the automated method generator embedded in the Analyst software (Sciex). Protein identification and quantification was conducted using Peak View software and the previously generated protein library.

**Immunohistochemistry & Immunofluorescence**

Five-micrometer thick paraffin sections were deparaffinized in xylene, rehydrated in a series of graded methanol dilutions (100% - 95% - 70% - 50%) and washed in phosphate saline buffer (PBS) with 0.25% Triton X-100 (VWR Chemicals, Randor, PA, USA; cat. no.: 28817.295). The endogenous peroxidase activity was blocked with 10% hydrogen peroxide in methanol (Sigma Aldrich, St. Louis, MI, USA; cat. no.: 216763) for 30 min. Antigen retrieval was conducted using AR6 buffer (Perkin Elmer, Waltham, MA, USA; cat. no.: AR600250) for 10 min in pressure cooker. The sections were blocked for 30 min in protein block serum-free solution (Agilent-Dako, Santa Clara, CA, USA; cat. no.: X0909) and incubated with the primary antibody at RT for 2h. The list of antibodies used in the present work is outlined in **Suppl. Table S1** below. Following this, the slides were washed three times in PBS for 5 min and then incubated for 30 min at RT with secondary antibody Histofine MAX PO Multi (Nichirei, Tokyo, Japan; cat. no. 414152F) for mouse and rabbit antibodies and Histofine MAX PO G (Nichirei Bio, cat. no. 414162F) for antibodies of goat origin. Subsequently, the sections were washed three times for 5 min in PBS and then stained with 3,3'-diaminobenzidine (DAB). The latter solution was made by adding 10µL of DAB Chromogen to 1 mL of DAB Substrate Buffer (Agilent-Dako, cat. no.: GV800). The slides were counter-stained in hematoxylin (Sigma Aldrich, cat. no.: MHS32) and mounted with Eukitt (Orsatech GmbH, Bobingen, Germany).

For immunofluorescence, tissue sections were prepared as described above with exception of primary antibody incubation that was conducted at 4°C and over night and the staining that has been performed using Opal system (Perkin Elmer, cat. no.: NEL810001KT). Following the primary antibody incubation, the slides were incubated with the corresponding secondary antibody as described above. The slides were then incubated with 100µL staining solution prepared from 2µL Opal dye and 98µL Amplifying Buffer. Following 10 min incubation, the slides were washed three times for 5 min in PBS and then subjected to microwave-assisted antibody removal. Slides were immersed in AR6 buffer and were treated in microwave for 15 min, maintaining the heat close to boiling point. After cooling and a wash in PBS buffer for 5 min, the tissues were re-blocked with for 30 min in protein block serum-free solution at RT. Tissues were then incubated with the next primary antibody and the staining procedure was repeated as described above using the following Opal dyes: 520, 570, 620 and 690. Finally, slides were mounted using VECTASHIELD® Antifade Mounting Medium with DAPI (Vector, Burlingame, USA).

**ifLR-score calculation and usage**

The determination of the percentage of receptor-expressing cells that are surrounded by sufficient ligand fluorescence relies on the computation of an immunofluorescence ligand-receptor score (ifLR-score) and the definition of a threshold above which the interaction is considered positive. Taking the PD-1/PD-L1 interaction as an example, we first determined the average diameters of PD-1+ and PD-L1+ cells independently (**Suppl. Fig. S1A**). This allowed us to define a crown-shaped area around each PD-1+ cell. The receptor abundance $R$ is estimated by the average PD-1 fluorescence inside the inner disc that is centered on the PD-1+ cell and has corresponding diameter. The ligand abundance $L$ is estimated in the crown that has a width equal to half the diameter of a PD-L1+ cell. $L$ represents the amount of ligands close enough to the PD-1+ cell to engage inhibition. Empirically, we define

$$\text{ifLR-score} = L^{1/3}R^{1/2}/(M + L^{1/3}R^{1/2}),$$

where $L$ and $R$ are as above, $M$ is the average of the average intensity over the whole ligand image and the average intensity over the whole receptor image (each label results in a separate gray-scale image). The fractional powers account for the ligand and the receptor to reside in a 3-, respectively 2-, dimensional space. $M$ represents the background signal intensity and its role is to regularize the ifLR-score to obtain values between 0 and 1. Analysis of the PD-1/PD-L1 interaction in 3 SDC allowed us to plot ifLR-score value distribution (**Suppl. Fig. S1B**). It is bimodal (or even trimodal for SDC22), with a fist mode corresponding to random signals (low values) followed by a rightmost mode corresponding to overlapping ligand and receptor fluorescence. We empirically set a conservative threshold at 0.4. That is, each PD-1+ cell with ifLR-score > 0.4 is considered in positive interaction with adjacent PD-L1+ cell(s), otherwise the interaction is deemed negative.

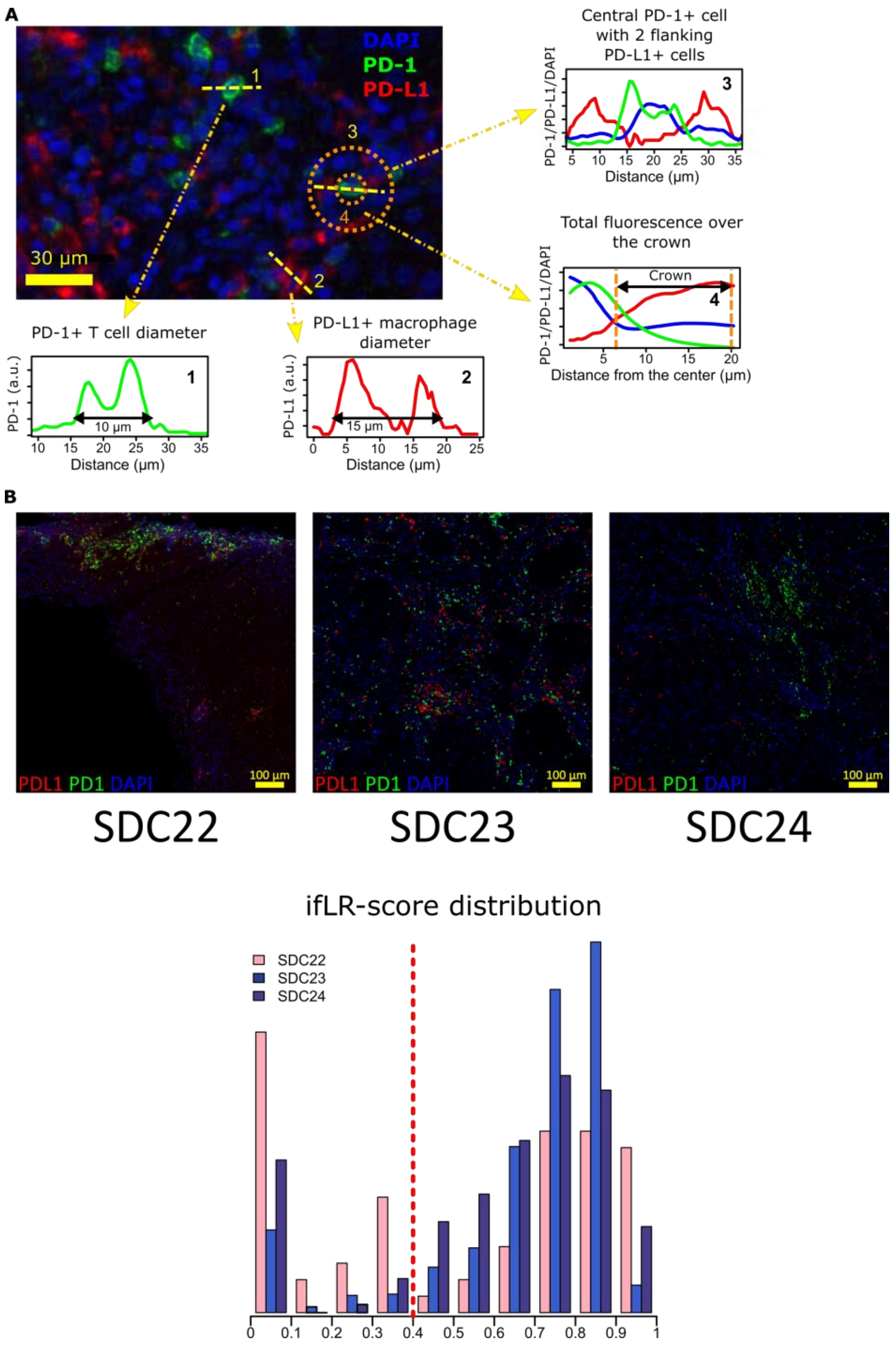


**Supplementary Figure S1. .** Principle of detecting ligand positive and receptor positive cell interactions exemplified by PD-1/PD-L1. **A.** Average PD-1+ cell diameter was estimated from 20 cells along a crossing axis (1). We note the increased green signal at the membrane and its decrease at the center of the cell (nucleus). Same operation for PD-L1+ cells (read, 2). Over a PD-1+ cell with flanking, adjacent PD-L1+ cells, we note the coherent signals with DAPI in blue (3). To compute the ifLR-score, we use the green average signal inside the inner circle of the crown (PD-1+ cell diameter) and the red average signal in the crown that has a width equal to half a PD-L1+ cell diameter. **B.** SDC IF images and ifLR-score distributions with the threshold as vertical dashed red line.

**LR-score (transcriptomics)**

We also defined a ligand-receptor score meant to assess co-expression in transcriptomics as a proxy for potential true interaction in the sample. The empirical formula is similar to above:

$$\text{LR-score} = l^{1/3} r^{1/2} / (\mu + l^{1/3} r^{1/2}),$$

where $l$ is the ligand read count in $\log_{10}$ (ligand transcript expression), $r$ the receptor read count in $\log_{10}$, and $\mu$ the average $\log_{10}$ read count over all the genes and all the SDC transcriptomes. See above for the fractional powers and $\mu$ roles.

#### Additional Tables and Figures

**Supplementary Table S2.** References and dilutions of primary antibodies used for IHC and IF.

| Target | Manufacturer | Cat. No. | Dilution IHC | Dilution IF |
|---|---|---|---|---|
| CD3 | Dako | GA503 | undiluted | 1/3 |
| CD8 | Roche | 5937248001 | undiluted | N/A |
| CD68 | Dako | M0814 | 1/5000 | 1/1500 |
| PD-L1 | Dako | M3653 | undiluted | 1/3 |
| CD163 | Roche | 7604437 | undiluted | 1/3 |
| α-SMA | Dako | M0851 | 1/500 | N/A |
| TIM3 | Cell Signalling Technology | 45208 | N/A | 1/150 |
| PD-1 | Cell Signalling Technology | 86163 | N/A | 1/300 |
| Galectin-9 | Cell Signalling Technology | 54330 | N/A | 1/2500 |
| Pan-Cytokeratin | Dako | GA053 | N/A | 1/3 |
| CTLA-4 | Abcam | ab227709 | undiluted | 1/50 |

**Supplementary Table S3.** Interactions added to Reactome binary interactions as retrieved from PathwayCommons. Information taken from UniprotKB/Swissprot.

| Receptor | Interactor | Receptor | Interactor | Receptor | Interactor |
|---|---|---|---|---|---|
| HAVCR2 | LCK | TNFRSF4 | TRAF2 | TNFRSF18 | TRAF2 |
| HAVCR2 | PLCG | TNFRSF4 | TRAF3 | TNFRSF18 | TRAF3 |
| HAVCR2 | VAV1 | TNFRSF4 | TRAF5 | TNFRSF18 | SIVA1 |
| HAVCR2 | AKT1 | TNFRSF8 | TRAF1 | TNFRSF25 | TNFRSF1 |
| HAVCR2 | AKT2 | TNFRSF8 | TRAF2 | TNFRSF25 | TRADD |
| HAVCR2 | LCP2 | TNFRSF8 | TRAF3 | TNFRSF25 | BAG4 |
| HAVCR2 | ZAP70 | TNFRSF8 | TRAF5 | | |
| HAVCR2 | SYK | TNFRSF9 | TRAF1 | | |
| HAVCR2 | PIK3R1 | TNFRSF9 | TRAF2 | | |
| HAVCR2 | FYN | TNFRSF9 | TRAF3 | | |
| HAVCR2 | SH3BP2 | TNFRSF9 | LRR1 | | |
| HAVCR2 | SH2D2A | TNFRSF18 | TRAF1 | | |

**Supplementary Table S4.** GO terms and Reactome pathways regulated in transcriptomics.

See external file.

**Supplementary table S5.** GO terms and Reactome pathways regulated in proteomics.

See external file.

**Supplementary Table S6.** IHC analysis.

See external file.

**Supplementary Table S7.** Confident LR pairs (179).

See external file.

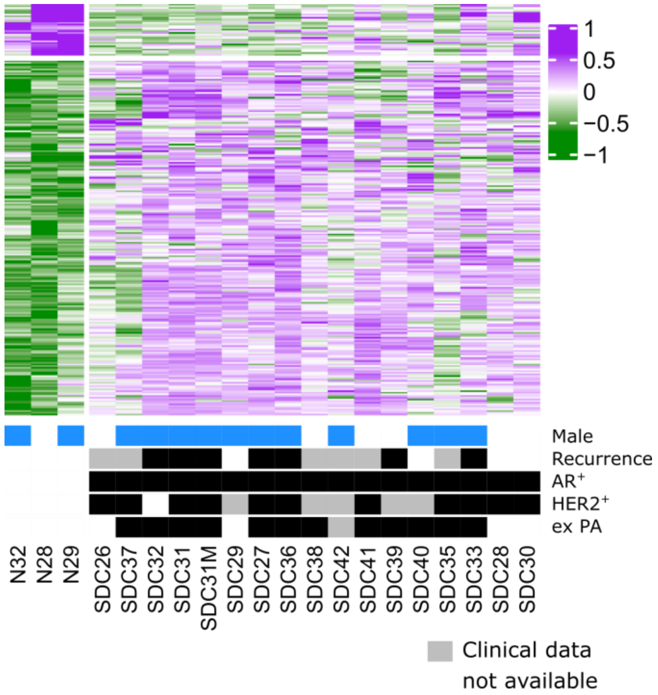


**Supplementary Figure S2.** Differentially expressed proteins. (Missing recurrence data are due to limited follow up time for recently enrolled patients).

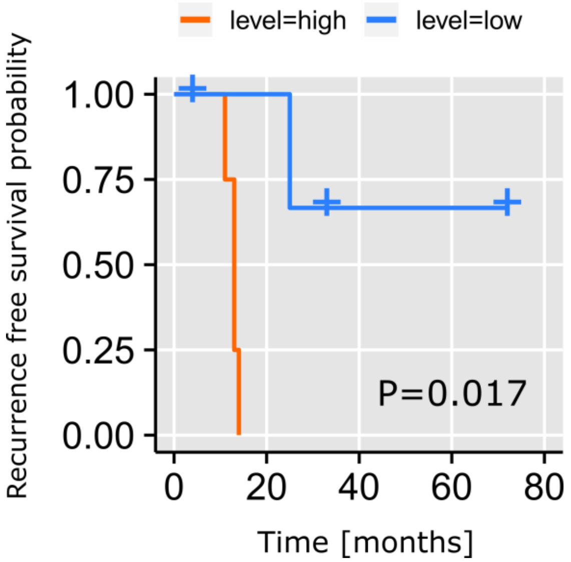


**Supplementary Figure S3.** Recurrence free survival our cohort 1 with respect to *IFNG* expression level (Kaplan-Meier curves, log-rank test, n=8, high=above median, low=below median).

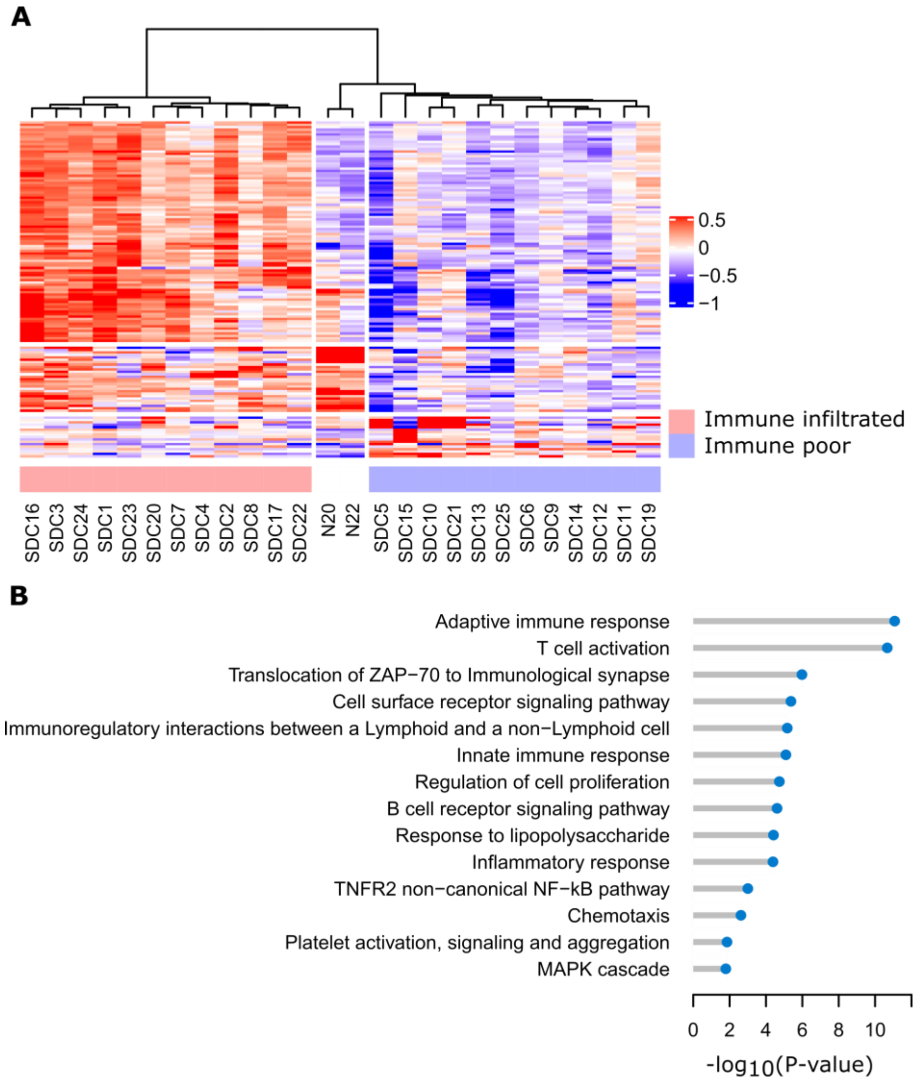


**Supplementary Figure S4.** Differentially expressed genes between immune infiltrated and poor SDC. (**A**) The comparison selected 135 significantly regulated genes (FDR<0.01, $log_2$-FC>2 in absolute value, average read count>20), which segregate the two sample clusters perfectly (plus normal samples for reference). (**B**) Main GOBP terms and Reactome pathways significantly enriched (hypergeometric test, FDR<0.05, minimum 5 regulated genes in the GO term or the Reactome pathway).

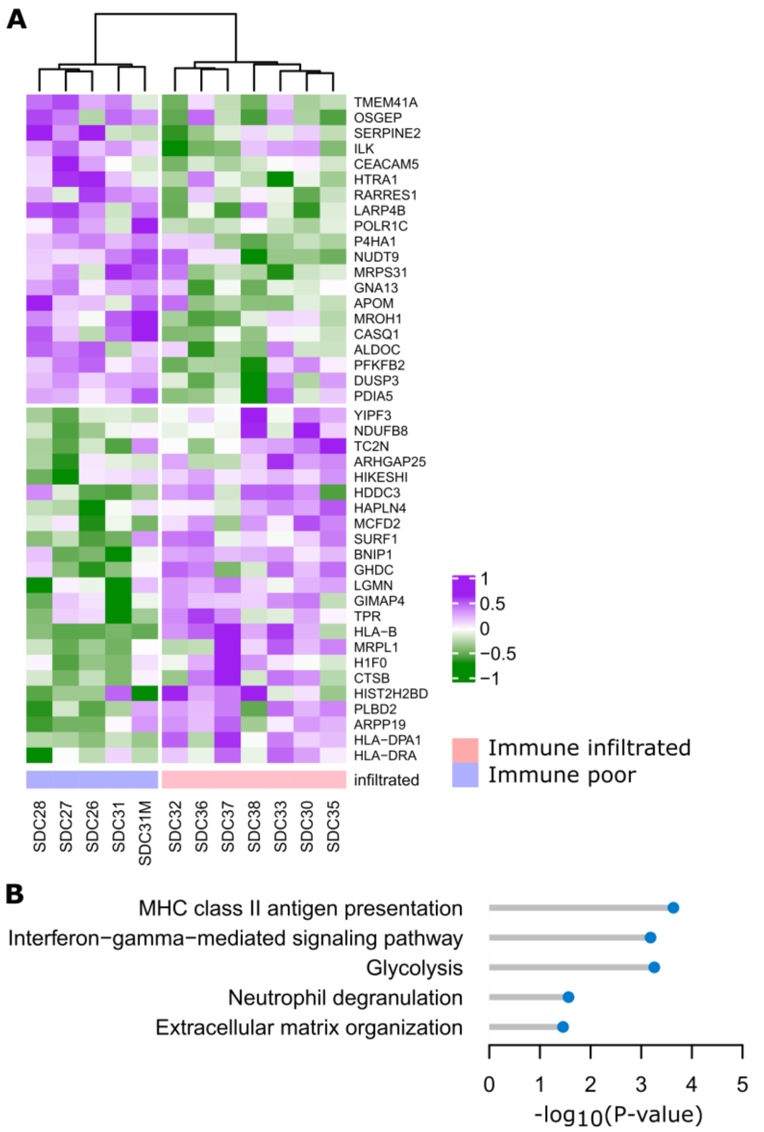


**Supplementary Figure S5.** Differentially expressed proteins between immune infiltrated and poor SDC. (**A**) The comparison selected 43 significantly regulated genes (P<0.05, FC>1.5, average MS signal >3), which segregate the two sample clusters perfectly (plus normal samples for reference). Proteomics data were available for 12/14 cohort 2 samples. (**B**) Main GOBP terms and Reactome pathways significantly enriched (hypergeometric test, FDR<0.05, minimum 3 regulated proteins in the GO term or the Reactome pathway).

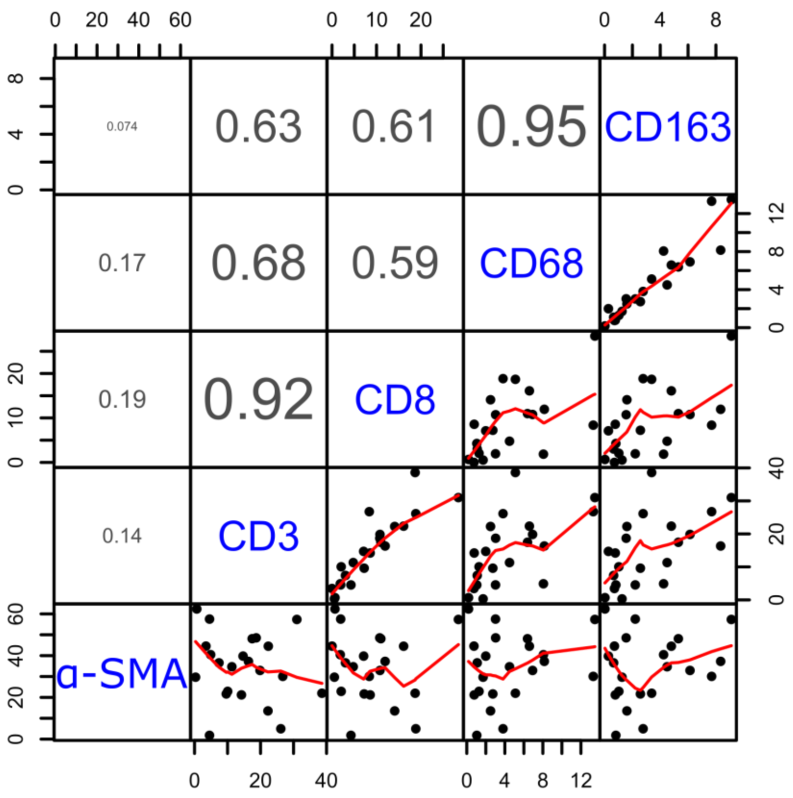


**Supplementary Figure S6.** Spearman correlations between CD3, CD8, CD68, CD163, and α-SMA levels. We note the quasi-absence of correlation with α-SMA indicating an immune infiltrate that does not depend on the desmoplastic stromal reaction level in SDCs (n=22).

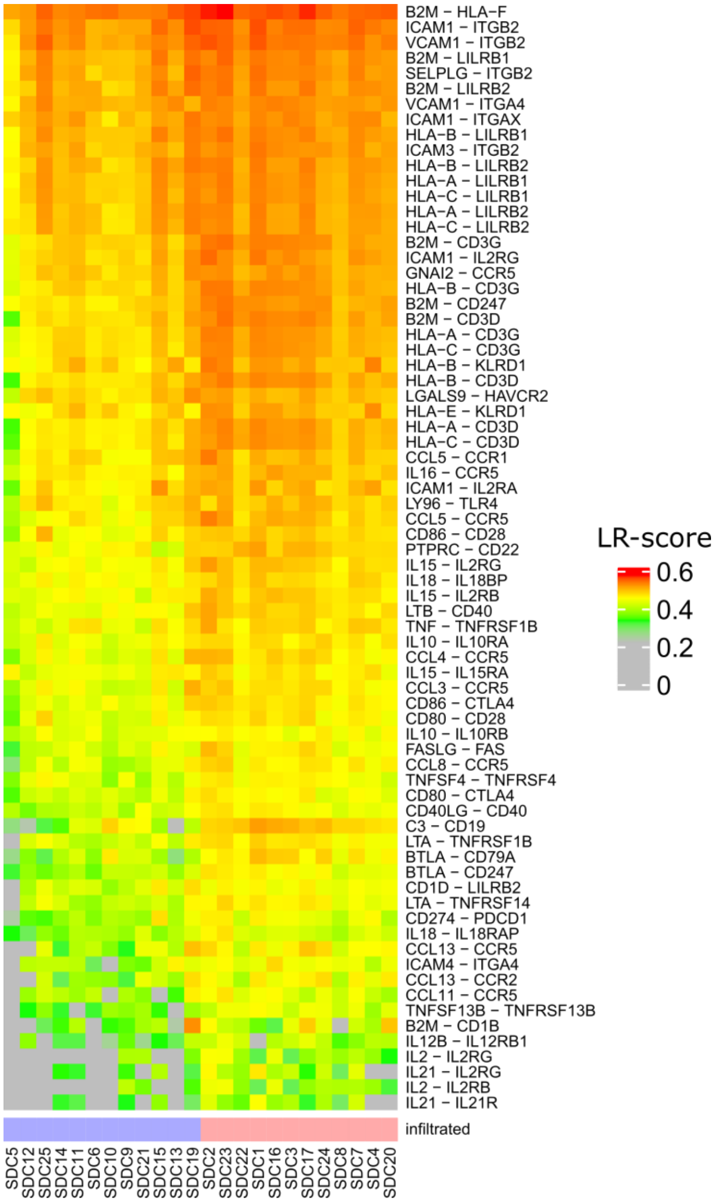


**Supplementary Figure S7.** Confidence LR pairs whose transcriptomic LR-score are correlated with immune infiltrate levels of SDC (Spearman r>0.6, immune infiltrate level defined as the average MCP-counter T cells, B cells and CD8+ cells signatures).

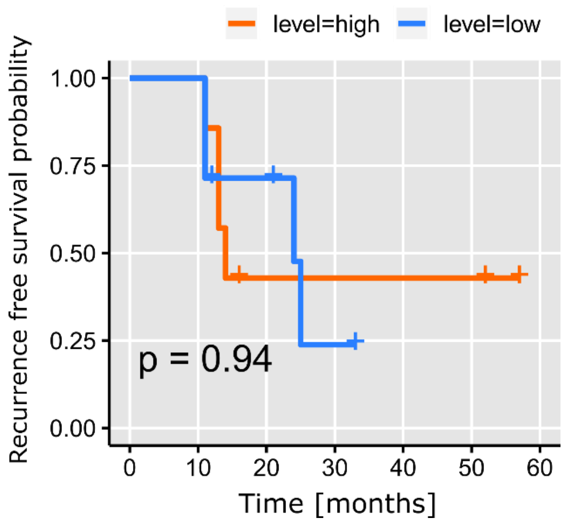


**Supplementary Figure S8.** Recurrence free survival among the immune infiltrated SDC with respect to the percentage of CD8+ positive cells. We note no significant association (Kaplan-Meier curves, log-rank test, n=14, high=above median, low=below median).

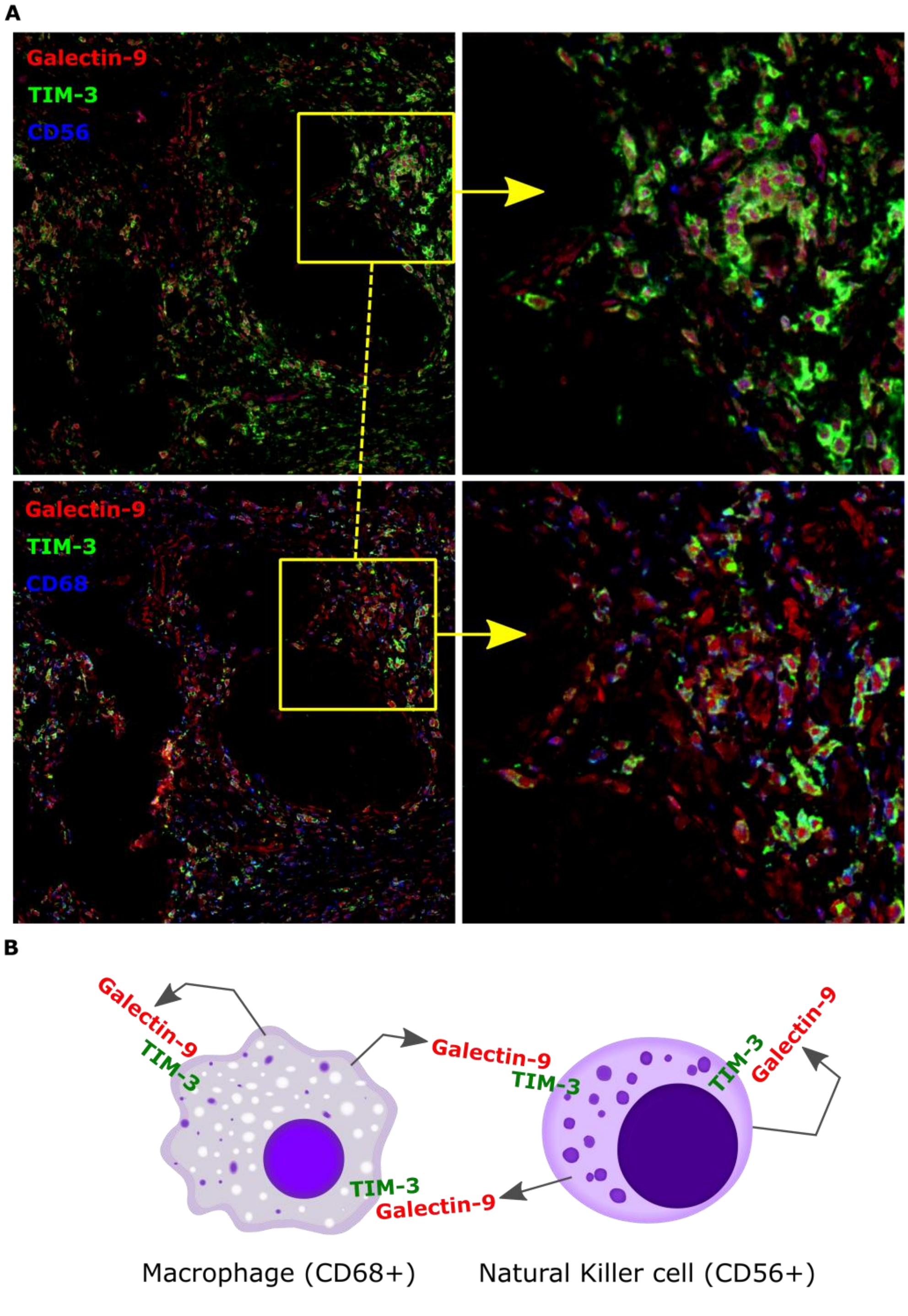


**Supplementary Figure S9.** Co-localization patterns of CD56, CD68, Galectin-9, and TIM-3 fluorescence. **A.** In adjacent FFPE block slides, we selected corresponding areas (yellow squares, left side). In both cases, we see individual cells positives for the markers (right side, CD68, Galectin-9, TIM-3 or CD56, Galectin-9, TIM-3) indicative of autocrine inhibition. Since the slides were adjacent and we selected the same areas, it additionally indicates simultaneous paracrine inhibition between macrophages and NK cells. **B.** Schema of dual auto- and paracrine inhibition. (Cell cartoon source: commons.wikimedia.org).
